# An early phase of instructive plasticity in visual cortex before the typical onset of sensory experience

**DOI:** 10.1101/635870

**Authors:** Arani Roy, Shen Wang, Benyamin Meschede-Krasa, Jordan Breffle, Stephen D. Van Hooser

**Author notes:** equal contribution (Arani Roy, Shen Wang). **Corresponding author:** Stephen D. Van Hooser, Brandeis University, 415 South St. MS008, Waltham, MA 02454 USA. **Contributions** AR and SDV designed experiments, AR, SW, and BMK performed data collection, AR, JB, and SDV performed data analysis, AR and SDV wrote the paper with input from all authors.

## Abstract

While early experience with moving stimuli is necessary for the development of direction selectivity in visual cortex of carnivores, it is unclear whether experience exerts a permissive or instructive influence. To test if the specific parameters of the experienced stimuli could instructively sculpt the emergent responses, visually naive ferrets were exposed to several hours of experience with unusual spatiotemporal patterns. In the most immature ferrets, cortical neurons developed selectivity to these patterns, indicating an instructive influence. In animals that were 1-10 days more mature, exposure to the same patterns led to a developmentally-typical increase in direction selectivity. We conclude that visual development progresses via an early phase of instructive plasticity, when the specific patterns of neural activity shape the specific parameters of the emerging response properties, followed by a late phase of permissive maturation, when sensory-driven activity merely serves to enhance the response properties already seeded in cortical circuits.

## Introduction

In the developing visual system, molecular cues^1,2^ and early spontaneous activity^3–7^ lay down the foundation of initial circuitry that exhibits many of the properties that are found in the mature animal, including retinotopic organization and orientation selectivity^8–12^. During a subsequent phase of experience-dependent development, visually-driven activity further shapes these response properties, providing enhanced cortical acuity^13^, binocular matching of inputs from the 2 eyes^14^, and, in carnivores and primates, the emergence of direction-of-motion selectivity^15,16^. It is of particular interest to understand how early visual activity interacts with, and alters, the immature circuit. Do the circuit connections established before the onset of experience commit cortex to a developmental path with pre-destined response properties, such that subsequent sensory experience merely permits maturation of these pre-seeded properties? Or is the cortical circuit malleable enough so that the particular patterns of visually-driven activity can instructively sculpt the responses according to the quality of the specific stimuli experienced?

Direction selectivity – a preference for stimulus movement in 1 direction as opposed to all others – typically develops in ferret visual cortex over a period of 1-2 weeks after eye opening through a process that requires visual experience^15,17,18^, and does not form in dark-reared^15^ or strobe-reared^19–22^ animals. Direction selectivity can also be rapidly induced in the laboratory by providing an anesthetized ferret kit with 3-9 hours of experience with drifting gratings^17,18,23,24^. While exposure to such smooth spatiotemporal motion increases direction selectivity, many parameters of direction tuning are invariant to the specific parameters of the gratings used for visual stimulation. For example, orientation selectivity is barely malleable during motion exposure: only columns whose orientation preference match the provided stimulus exhibit increases in direction selectivity^17^, and the orientation preferences of neurons that initially prefer other orientations are changed only very slightly^17^. Direction angle preference is also relatively unchangeable: stimulation with gratings that move in only one direction cause a dramatic increase in direction selectivity for cells whose initial biases match the stimulated direction, but do not cause an increase in selectivity for cells whose initial biases match the opposite direction^23^. Speed / temporal frequency tuning is also invariant: stimulation with either slow or fast moving stimuli causes an increase in direction selectivity, but does not alter tuning for speed / temporal frequency^24^. These results suggest that experience with drifting gratings fails to modify many of the parameters of direction tuning (orientation/direction preference angle, speed/ temporal frequency etc.), thereby implying that visual experience is only necessary to permissively increase selectivity and acuity of the tuning.

While the above results suggest a limit to the extent to which the experienced stimulus can shape cortical tuning properties, no experiment to date has directly tested if the nascent visual cortex can be induced to develop selective responses to irregular spatiotemporal patterns, which would be a strong test of whether selectivity is instructed by activity. In all visual motion stimulation experiments to date, young ferrets were exposed to smoothly moving gratings, in which an oriented grating is moved along a smoothly progressing sequence of spatial phases in time. According to the spatiotemporal receptive fields of neurons in the typically-developed visual cortex, such stimuli are ideally suited for driving cortical neurons^25–27^. In addition, the vast majority of ferret kits examined in prior studies already had visual experience for 1-3 days at the time of each experiment, making it difficult to rigorously assess if activity before or around the time of natural eye opening could instructively modify the cortex.

To address these issues, we directly manipulated early visual experience by prematurely opening the eyes of young ferrets and exposing them to grating stimuli that were animated with scrambled spatiotemporal phase sequences. We reasoned that if the patterns of early activity in visual circuits were instructive, then we should be able to induce increased responses to these phase-scrambled grating stimuli through repeated visual exposure. On the other hand, if the cortical circuitry were already committed to developing selectivity for smooth motion, then providing phase-scrambled stimulation should merely increase direction selectivity.

We found evidence for a transition of the influence of early activity in the visual cortex – from instructive to permissive – that occurred around the time of natural eye opening. When the eyes were opened prematurely, or if the state of the cortex was very immature as assessed by levels of orientation selectivity, animals developed increased selectivity to the artificial phase-scrambled stimulus that was experienced.

Animals that were slightly more mature did not acquire increased selectivity to the phase-scrambled patterns but instead exhibited a developmentally-typical increase in direction selectivity, consistent with a permissive influence of visual stimulation. These data provide evidence that the early activity in visual cortex that occurs before and at eye opening – which includes spontaneous activity^3–5^, low resolution visual stimulation through the closed lids^28,29^, and higher resolution vision through the slowly opening eyes – provides an instructive signal for neural circuit construction. Later activity, after the normal onset of visual experience, is necessary for the maturation of direction selectivity, but only in a permissive manner.

## Results

Neurons in carnivore primary visual cortex respond strongly to oriented gratings moving in one direction following a smoothly progressing sequence of spatiotemporal phases. We wanted to test if early exposure to gratings moving with irregular spatiotemporal patterns could modify the cortical circuitry and induce neurons to respond selectively to irregular motion. For this purpose, we designed a stimulus family of gratings moving with scrambled spatiotemporal phase sequences. To create such phase-scrambled visual stimuli, we varied the typical oriented grating stimuli that drive the cortex well. We discretized grating phase into 8 steps (**Figure 1**), defined forward (F) and backward (B) stimuli as phase sequences [1 2 3 4 5 6 7 8] and [8 7 6 5 4 3 2 1], respectively, and approximated a viewing temporal frequency of 2 Hz by showing each phase for 1/(8*2Hz) = 0.0625 s. We quantitatively analyzed the set of possible 5040 unique sequences (see Supplementary methods; **Figure 1C, Supplementary Figure S1, S2**), and evaluated their degree of similarity to smooth motion. Subsequently, we chose for experiments a family of 10 sequences, containing a mixture of low and intermediate levels of similarity to smooth motion: forward motion (F), backward motion (B), 6 sequences that exhibited varying degrees of correlation with forward and backward motion (scrambled: *S1-S6*), and counterphase stimuli at 2 spatial phases labeled *CP1* and *CP2*, respectively (**Figure 1ABCD; Supplementary Videos 1-10**). Stimuli *S1-S6* contain spatiotemporal energy at multiple spatial and temporal frequencies (**Supplementary Figure S2**), while stimuli *F*, *B*, *CP1*, and *CP2* contain energy around a single spatial and temporal frequency. *S4* and *S6* were chosen for visual stimulation due to their very low correlation with smooth motion, while all 10 sequences were used to test responses before and after visual stimulation.

**Figure 1.**
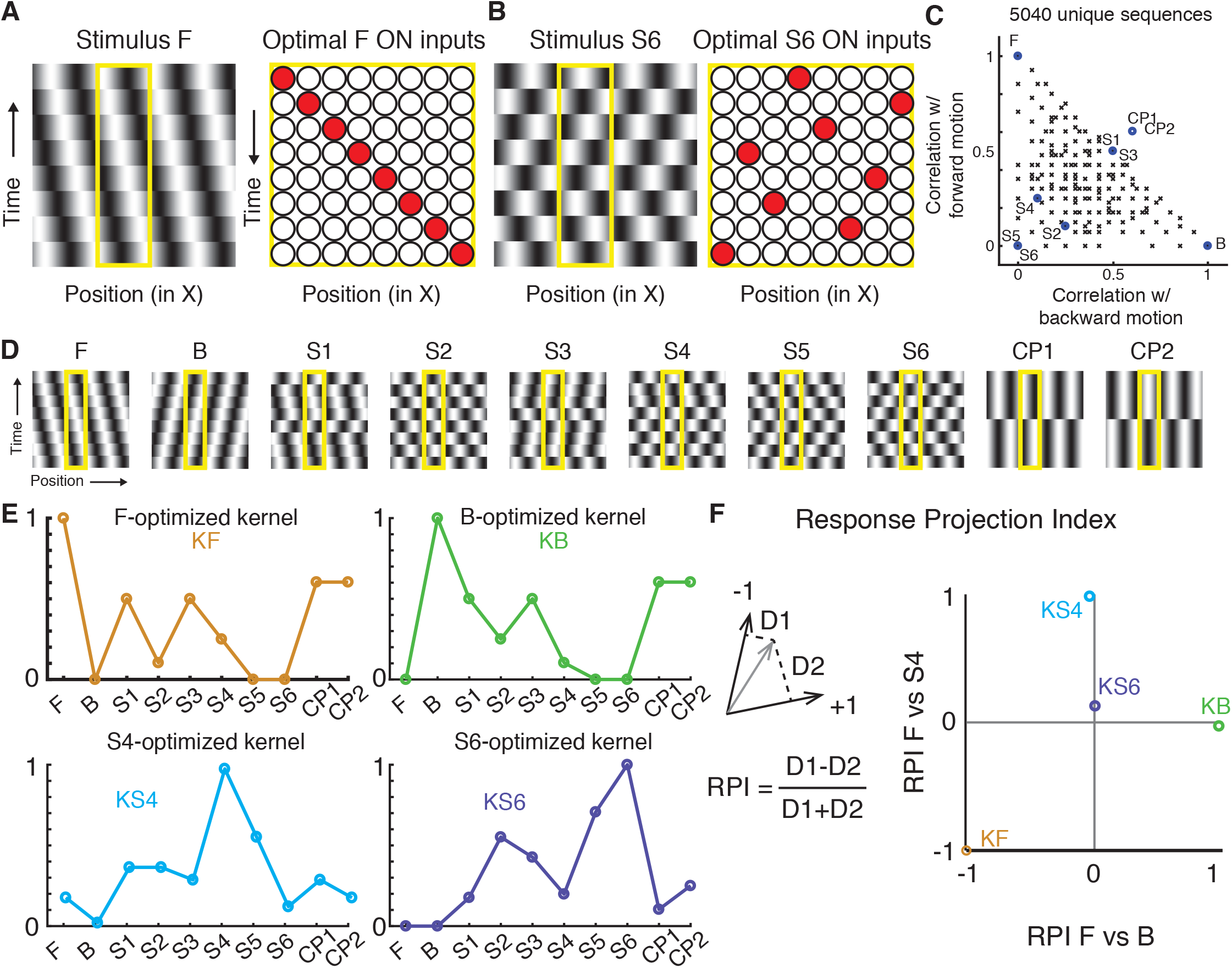
Design of a spatiotemporal stimulus family. **A)** Left: X-T (space-time) view of a vertical sinusoidal grating shifting to the left at each phase step, termed forward stimulus (F). Each strip represents a video frame. The phase progression of this grating is denoted as [1 2 3 4 5 6 7 8]. Yellow box indicates hypothetical location of a receptive field. Right: Hypothetical X-T inputs (ON only) that depicts positions and latencies of inputs that would drive an optimal response to stimulus F. **B)** Left: X-T view of a vertical sinusoidal grating advancing with scrambled phase steps ([8 3 6 2 7 4 1 5]), termed stimulus S6. Right: Hypothetical X-T inputs (ON only) that depicts positions and latencies of inputs that would drive an optimal response to stimulus S6. **C)** Plot of the best-aligned correlation with forward (F) and backward (B) motion for all 5040 possible unique phase sequences (black x) and our selections for stimuli marked as blue circles. F and B represent the forward and backward smooth motions, respectively; S1-S6 are phase-scrambled stimuli that deviate substantially from forward or backward motion; CP1 and CP2 are counterphase stimuli at 2Hz (phase progression [1 1 1 1 5 5 5 5] and [3 3 3 3 7 7 7 7], respectively). **D)** Video frame strips of all 10 chosen sequences. **E)** Responses of hypothetical cells with input kernels optimized for indicated stimuli. A cell optimized for F *(KF*, orange) responds strongly to F, but not to B, S5, or S6. A cell optimized for B *(KB*, green) responds strongly to B but not F, S5, or S6. Cells optimized for either S4 *(KS4*, cyan) or S6 *(KS6*, purple) respond weakly to F and B and relatively poorly to each other’s optimal stimulus. **F)** Response Projection Index (RPI) indicates how well the tuning curve of a given cell matches that expected by a cell optimized for 2 particular stimuli. *Left*: Each cell’s normalized response curve in 10-dimensional space. The response is compared to the responses expected from hypothetical reference neurons that are optimized for 2 given stimuli (such as F and B). The distance in vector space between the actual response (gray vector) and the vector line that defines the 2 reference neurons is calculated (D1 and D2), and an index is calculated RPI = (D1-D2)/(D1+D2). If the cell’s responses match that expected by a hypothetical neuron that is optimized for the first reference stimulus, then RPI is −1. If the cell’s responses match that expected by a hypothetical neuron that is optimized for the second stimulus, RPI is 1. *Right:* Scatter plot of RPI index values for kernels optimized for the particular stimuli indicated. X axis is RPI relative to F and B and Y axis is RPI relative to F and S4.

We developed 2 selectivity measures to quantify neural responses to this stimulus family – the response set for each neuron being 10-dimensional due to the inclusion of 10 stimuli in the experiment. The first measure, called the Selectivity Index *(SI)* for stimulus n, is equal to the response of the neuron to that stimulus divided by the sum of the responses to the chosen 10 stimuli. We also developed a second measure called the Response Projection Index *(RPI)*, which considered the fact that *F…CP2* are correlated with one another to varying degrees. We can imagine linear receptive field kernels *(KX)* that would give a maximal response to a stimulus (*X*), as shown in **Figure 1E**, and we can compute the responses of these kernels to each of the 10 stimuli. The RPI describes how close the response of a measured neuron, in 10-dimensional response space, is to the responses that would be expected from an ideal kernel *(KX)* relative to another ideal kernel *(KY)* (**Figure 1F**). A neuron that gives responses identical to *KF* would have an *RPI(KF vs KB)* value of −1, whereas a neuron that gives responses that are identical to *KB* would have an *RPI(KF vs KB)* value of+1. Stimulus *S6* is uncorrelated with *F* and B, and *KS6* has an *RPI(KF vs KB)* value of 0 (**Figure 1f**).

We prepared young ferrets for 2-photon imaging of virally-expressed GCaMP6s in visual cortex^30^ (see Supplementary Methods). We began each experiment by measuring responses to drifting gratings in order to assess the initial orientation and direction tuning. A well-represented orientation preference was selected to be the orientation angle for stimuli *[F…CP2]*, and responses to these stimuli were measured. Initial responses were measured at a single depth in order to limit stimulation that could alter the receptive fields. Next, stimulus *S4* or *S6* (stimuli that were poorly correlated with *F* and B) was selected for 6 hours of prolonged visual stimulation^17,23,24^; in previous studies, 3-6 hours of visual stimulation was sufficient to drive substantial increases in direction selectivity, and animals remained physiologically robust over this time. Finally, responses to stimuli *[F…CP2]* and traditional orientation and direction tuning were re-assessed using sinusoidal gratings. In a few experiments, we were able to track some of the same cells over time, but in most experiments cells expressing GCaMP6s were dark when unstimulated, and exact alignment of imaging fields was not performed.

Example responses from a ferret (age P30) whose eyes were opened prematurely are plotted in **Figure 2**. After 6 hours of exposure to stimulus *S4*, there is a clear enhancement of the response of the imaging field to stimulus *S4* (**Figure 2A-D, H-J**). To characterize the degree to which neural responses were similar to that of a neuron that is perfectly selective to the trained stimulus *S4*, we computed the RPI for stimulus *F* vs *B* and *F* vs *S4*. There is a clear upward shift in *RPIF vs. S4*, indicating that neural responses are more selective for stimulus *S4* after exposure than before (**Figure 2H-J**). Despite the fact that the ferret received stimulation with the relatively broadband motion stimulus *S4*, traditional direction selectivity index values for this animal exhibited a decrease (**Figure 2D-F, KL**), which is opposite to what we would have expected if visual experience were only capable of inducing permissive changes^17,23,24^. Responses from another ferret (age P31) whose eyes were opened prematurely are shown in **Figure 3**. This animal was shown stimulus *S6* for 6 hours, and also exhibited an increase in response to stimulus *S6* (**Figure 3A-D, H-J**). This animal exhibited no significant change in direction selectivity (**Figure 3D-F, KL**), indicating that selectivity was reconfigured in a manner that, while closer to a hypothetical neuron that would respond to stimulus *S6*, did not change significantly in terms of direction selectivity.

**Figure 2.**
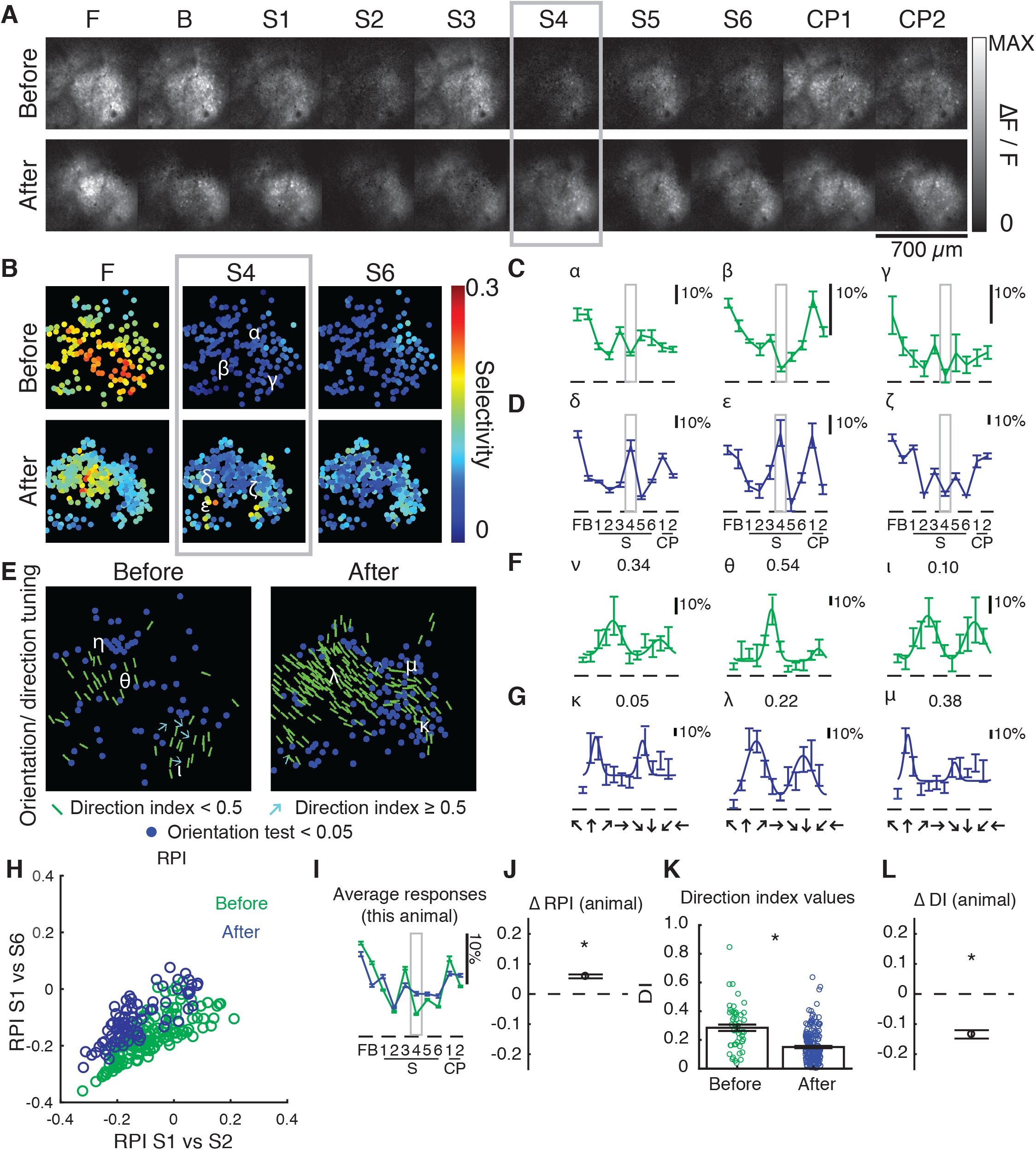
In a visually naïve ferret, 6 hours of experience with a phase-scrambled grating pattern caused an increase in selectivity for that pattern. **A)** Pixel map of GCaMP6s responses to a family of spatiotemporal stimuli before and after 6 hours of training with pattern S4, indicating a substantial increase in selectivity for *S4*. The animal’s eyes were opened prematurely on P30. **B)** Single cell Selectivity Index (SI) values for different stimuli (F – a phase-regular, unscrambled direction stimulus, and *S4/S6* – phase-scrambled stimuli). Selectivity for smooth motion (S1) decreases, while selectivity for stimulus *S4* increases in many cells. **C)** Responses to 3 example cells (indicated in B) before experience. Error bars are standard error of the mean (SEM) across trials. **D)** Responses to 3 example cells (indicated in B) after experience. Cells δ and ε exhibit strong responses to stimulus *S4*. **E)** Orientation and direction tuning in single cells before and after training. Blue dots represent visually-responsive cells that do not exhibit significant variation across stimuli moving in different directions; green bars represent orientation-selective but not strongly direction-selective cells (DI<0.5) and cyan arrows indicate strongly direction-selective cells (DI≥0.5). **F)** Direction tuning of 3 example cells (indicated in E) before experience. Numbers indicate direction index values. Error bars are SEM across trials. **G)** Direction tuning of 3 example cells (indicated in E) after experience. **H)** Response Projection Index (RPI) for *S1* vs *S2* (X axis) and *S1* vs *S4* (Y axis) for cells measured before (green) and after (blue) 6 hours of experience with *S4*. There is a substantial upward shift on the Y axis, indicating that cells exhibit responses that are more like a cell that is optimized to respond to *S4*. **I)** Grand average of responses before and after 6 hours exposure to *S4*. On average, there is an enhancement of the response to *S4*. Error bars are SEM across cells. **J)** Estimated difference in cell RPI *(S1* vs. *S4)* before (N=138 cells) and after (N=82 cells) experience (error bars are 95% confidence intervals), indicating a significant increase in selectivity to stimulus *S4*. * indicates that 95% confidence interval does not include 0. **K)** Direction index values of cells before (N=50 cells) and after (N=200 cells) exposure to *S4*. Direction index values decreased slightly after exposure to *S4*. Error bars are SEM across cells. * indicates that 95% confidence interval does not include 0 (see **L)**. **L)** Estimated difference in DI of cells before and after experience (error bars are 95% confidence intervals), indicating a significant decrease in DI with *S4* experience. * indicates that 95% confidence interval does not include 0.

**Figure 3.**
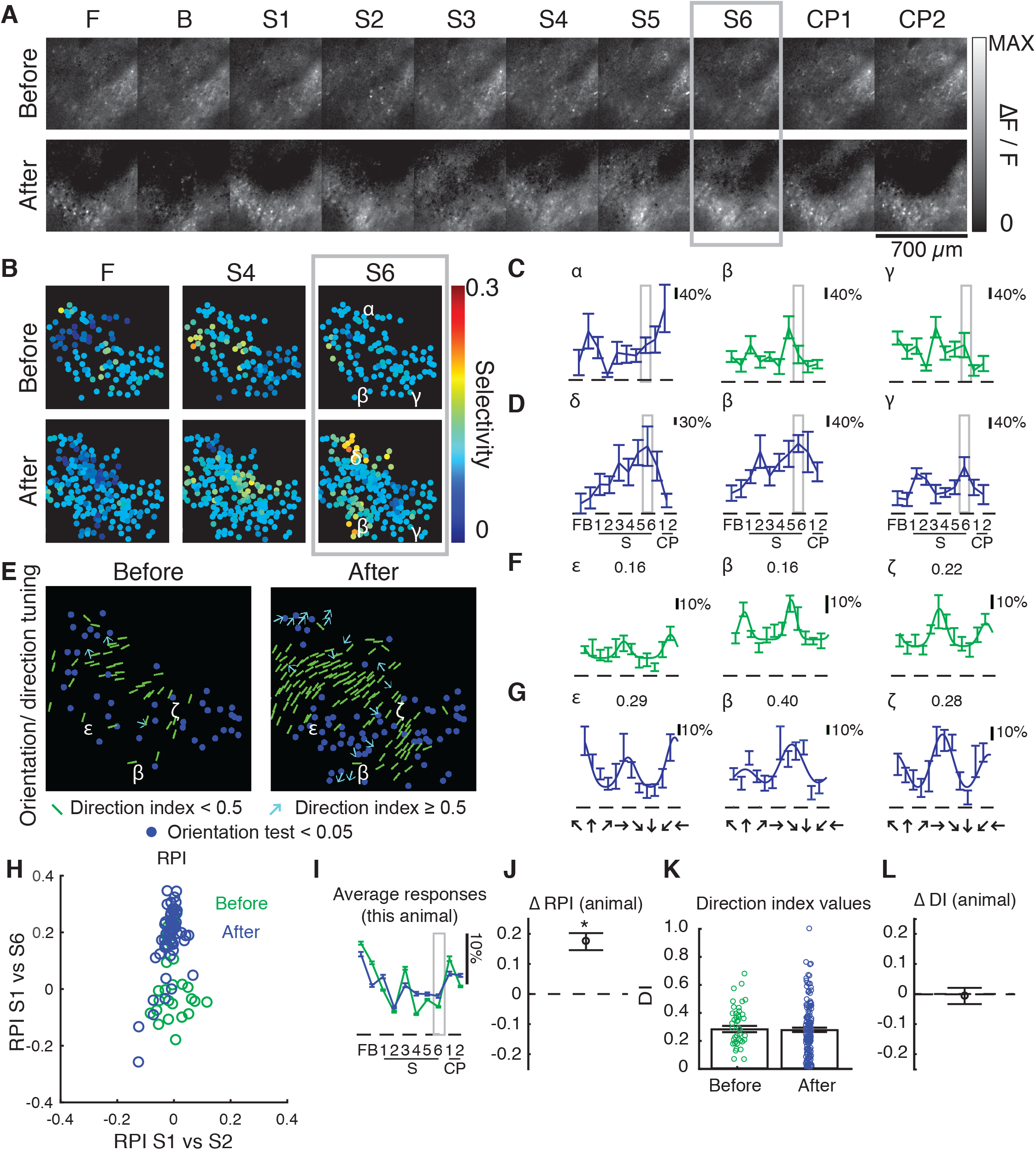
In a second visually naïve ferret, 6 hours of experience with a phase-scrambled grating pattern caused an increase in selectivity for that pattern. The eyes were opened prematurely on P31. Panels are as described in **Figure 2**, except that *S6* was used as the training stimulus. In this animal, some cells were tracked across time and Greek letters appear more than once. This animal exhibited increased selectivity to training stimulus S6, and no significant alterations to direction selectivity. N=25 cells before and N=62 cells after for measurements of F, B, S1-6, CP1-2, and N=41 cells before and N=145 cells after for direction tuning. Premature animals varied in the influence of phase-scrambled grating patterns on traditional direction selectivity.

Responses in slightly more mature ferrets resembled the example in **Figure 4**. This ferret (age P36, 3 days of natural visual experience) was also exposed to stimulus *S4* for 6 hours, following which there was no increase in selectivity to stimulus *S4* (**Figure 42A-D, H-J**). As a result, there was no upward shift in *RPIF vs. S4*, in fact there was a small but significant downward shift (**Figure 4H-J**). Instead, this animal exhibited an increase in direction selectivity index values, (**Figure 4D-F, KL**). A second example from an older animal (age 38, 5 days of visual experience) is shown in **Figure 5**. After 6 hours of exposure to stimulus S6, *RPIF vs. S6* in this animal showed no significant change (**Figure 5A-D, H-J**), but direction selectivity index was significantly increased (**Figure 5D-F, KL**). These results show that exposure to phase-scrambled stimuli in ferrets with some prior visual experience lead to enhancement of smooth-motion direction selectivity, which is consistent with a permissive role of visual experience for the development of direction selectivity.

**Figure 4.**
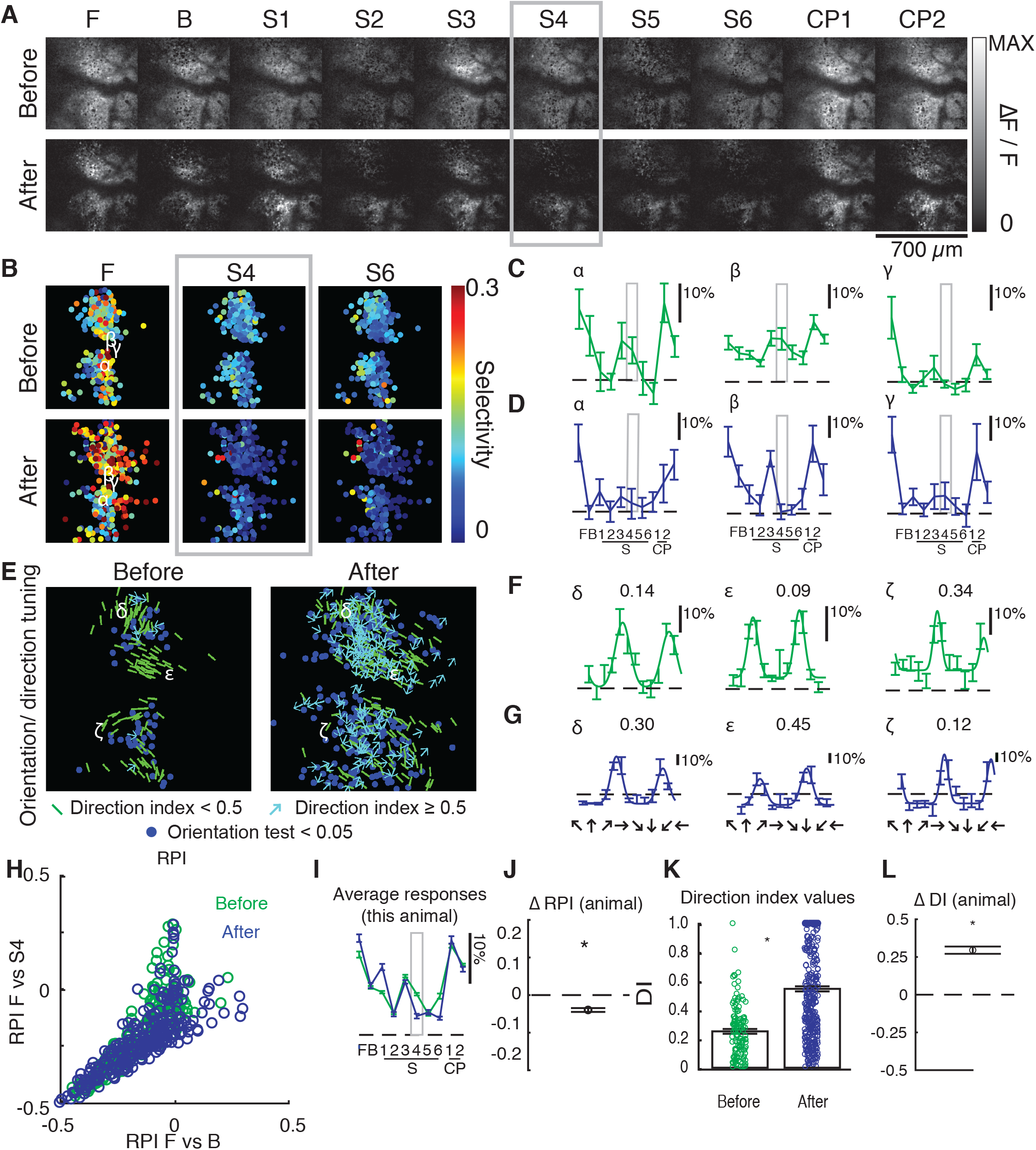
In a ferret with 3 days of visual experience, 6 hours of experience with a phase-scrambled grating pattern caused an increase in direction selectivity rather than selectivity for the phase-scrambled pattern. The eyes opened naturally on day P33, and the experiment was performed at P36. Panels are as described in **Figures 2 and 3**. *S4* was used as the training stimulus. In this animal, some cells were tracked across time and Greek letters appear more than once. N=170 cells before and N=267 cells after for measurements of F, B, S1-6, CP1-2, and N=124 cells before and N=371 cells after for direction tuning. This animal exhibited decreased increased selectivity to training stimulus *S4*, and exhibited a significant increase in direction selectivity.

**Figure 5.**
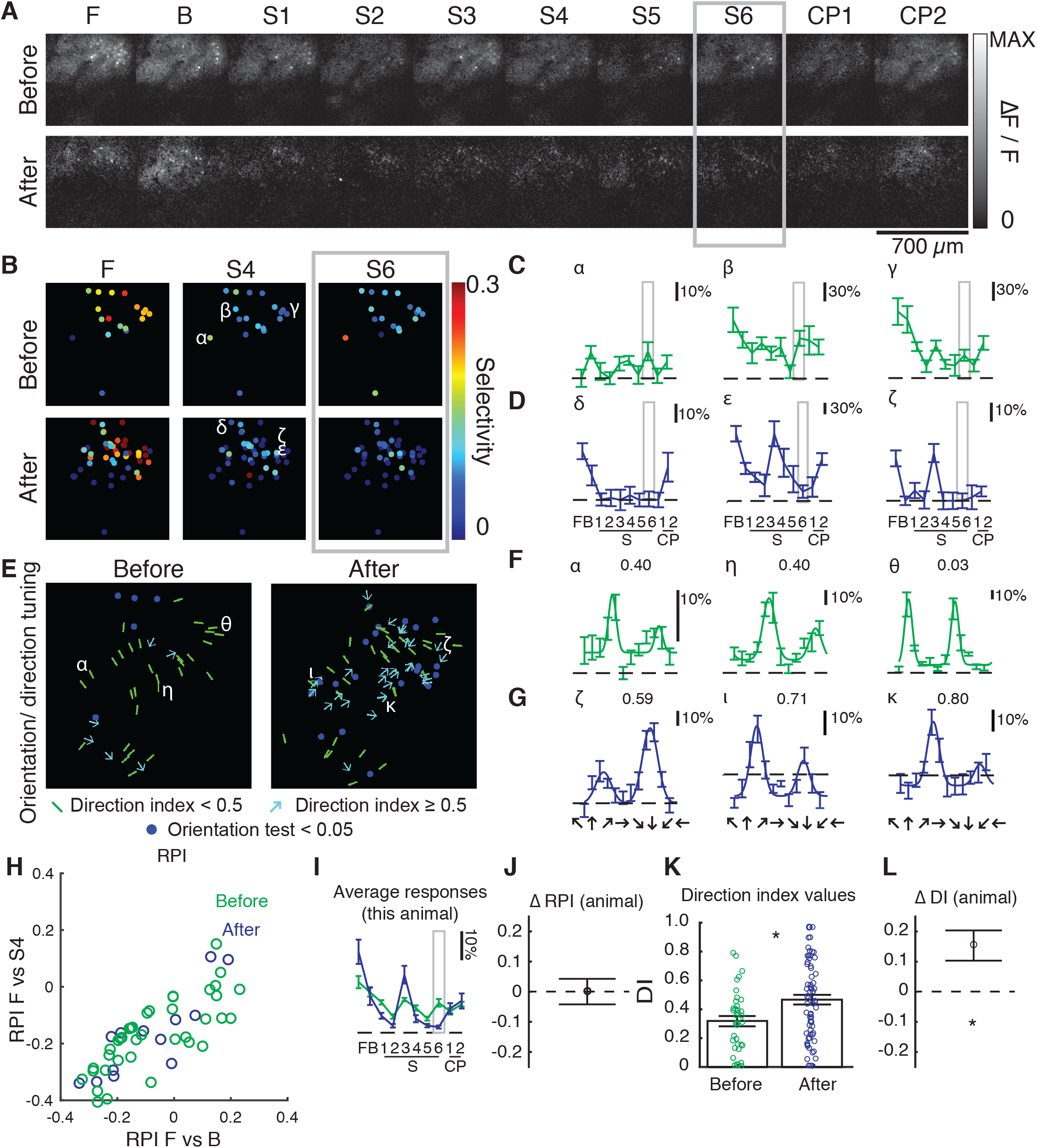
In a ferret with 5 days of visual experience, 6 hours of experience with a phase-scrambled grating pattern caused an increase in direction selectivity rather than selectivity for the phase-scrambled pattern. The eyes opened naturally on day P33, and the experiment was performed at P38. Panels are as described in **Figures 2–4**. *S6* was used as the training stimulus. In this animal, some cells were tracked across some trials and Greek letters appear more than once. N=13 cells before and N=41 cells after for measurements of F, B, S1-6, CP1-2, and N=36 cells before and N=67 cells after for direction tuning. This animal exhibited decreased increased selectivity to training stimulus *S6*, and exhibited a significant increase in direction selectivity.

Comparison of RPI and DI before and after visual stimulation for every ferret in the study is shown in **Supplementary Figure S3**. In all (4/4) ferrets with no visual experience there was a positive ΔRPI, suggesting increased selectivity for the training sequence following exposure (EO = 0, **Supplementary Figure S3ABCD**). In 3/4 of these same ferrets, there was a negative or zero ΔDI, suggesting no permissive increase in traditional direction selectivity. Taken together, these results imply that visual experience exerts an instructive role in young ferrets without prior visual experience. In contrast, in the majority (6/8) of young ferrets with several days of visual experience (EO 1-10; **Supplementary Figure S3EFGHIJKL**), ΔRPI was either slightly negative or not different from zero, suggesting lack of selectivity gain in favor of the training sequence. However, in most (7/8) of these slightly more experienced ferrets, there was a significantly positive ΔDI, suggesting increased direction selectivity, just as in animals from our 2008 paper (**Supplementary Figure S3N**). Taken together, these results imply that visual experience exerts a permissive role in young ferrets with longer (1-10 days) visual experience. A single older ferret imaged beyond the critical period for direction selectivity development (EO 20; **Supplementary Figure S3M**) showed a small decrease in both RPI and DI.

The above results suggest that the influence of early sensory experience-driven cortical activity on the nascent circuit undergoes a developmental transition – from instructive to permissive – just around the time of natural eye opening. We wanted to examine which out of a handful of parameters that all reflect different aspects of circuit maturity were correlated with increased selectivity to the phase-scrambled training stimulus or increased direction selectivity. We decided to concentrate on 2 parameters in particular, namely, eye-open status (EO) and initial orientation selectivity. Both these parameters increase with increasing age (**Figure 6AB**), but, because individual ferrets open eyes at different ages and mature at different rates, age alone is not a good predictor for level of experience or circuit maturity. Therefore, we concentrated on correlating ΔRPI and ΔDI with eye-open status (EO) and initial orientation selectivity. Ferrets whose eyes were opened prematurely (EO = 0) were highly likely (4/4) to exhibit increased selectivity to the phase-scrambled stimulus (**Figure 6CD**), but the hallmark of immaturity that best predicted increased selectivity to the phase-scrambled stimulus was the animal’s initial orientation selectivity index (OSI) value (**Figure 6E**). While orientation selectivity is evident at the time of eye opening, OSI values are relatively small in naïve animals and increase substantially over the first 1-2 weeks of visual experience^15,17^. Animals with weak initial OSI values showed large increases in selectivity for the phase-scrambled stimulus and lacked increases in direction selectivity, while animals with stronger initial OSI values (>0.3) generally lacked increases in selectivity for the phase-scrambled stimulus (6/8) and instead exhibited robust increases in direction selectivity (7/8) (**Figure 6EF**). We also analyzed the data by categorizing the ferrets into inexperienced (EO<1) and experienced (EO≥1) groups, or low (1-CV<0.3) and high (1-CV≥0.3) initial orientation selectivity groups. The results presented in **Figure 6GH** corroborate that RPI exhibits significantly larger increases in the inexperienced and low orientation selectivity index value groups compared to experienced or high initial orientation selectivity groups.

**Figure 6.**
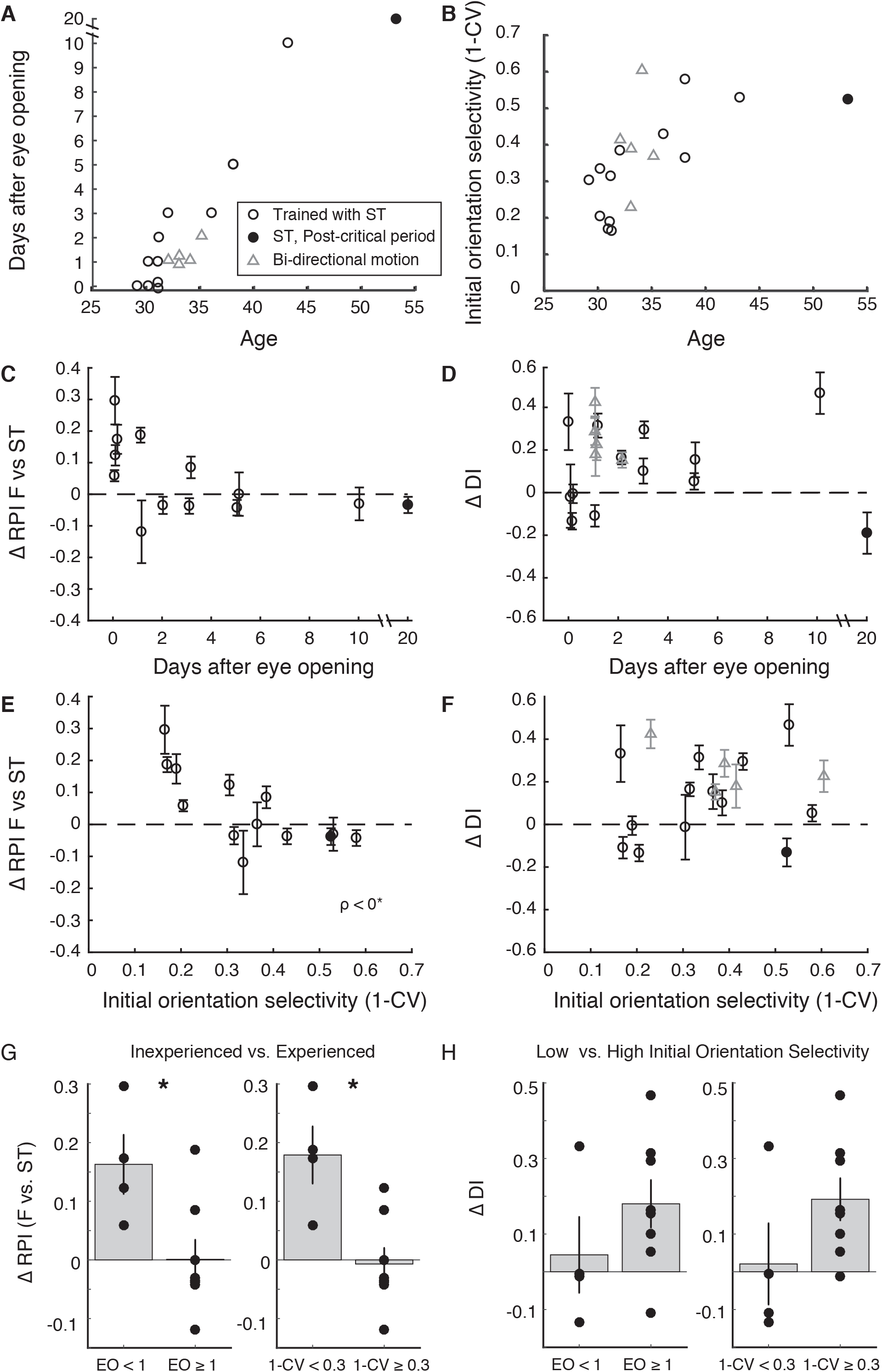
Relationship between changes in visual selectivity after phase-scrambled stimulus exposure and parameters that are related to animal maturity. **A)** Relationship between animal age and days after eye opening; ferrets exhibit a range of eye opening from P29 – P34. *ST* indicates animals that were trained with S4 or S6. Triangles indicate animals from Li/Van Hooser et al. (2008) that were trained with bidirectional moving stimuli. Filled circle is a single animal that was beyond the critical period for direction selectivity development (about 2 weeks after eye opening). **B)** Relationship between animal age and initial orientation selectivity as quantified by 1 minus the circular variance (CV). On average, orientation selectivity becomes stronger with age, but there is a range of initial selectivity in the youngest animals, which likely reflects the range of cortical maturity achieved. **C)** Difference in RPI for F vs the trained stimulus (denoted *ST;* S4 or S6, depending) (Y axis) before and after training (error bars are 95% confidence intervals) plotted against days of visual experience (days after eye opening). **D)** Same, but difference in direction index values is plotted. **E)** Difference in RPI vs. initial orientation selectivity that was measured at the beginning of the experiment (before training stimulus exposure). Animals with the lowest orientation selectivity exhibit strong changes in RPI and become more selective for the arbitrary training stimulus. p<0* indicates significant negative correlation (ρ<0.009, DF=12-2). **F)** Same, but for DI. **G)** *Left:* Changes in Response Projection Index (F vs. training stimulus ST) for animals whose eyes were opened by the experimenter (EO<1) and animals whose eyes had opened naturally before the experiment (EO≥1). * indicates significant difference by t-test (p<0.020241, DF = 12-2). *Right:* Changes in RPI (F vs.ST) for animals that exhibited low initial orientation selectivity (1-CV<0.3) and animals that exhibited higher initial orientation selectivity (1-CV≥0.3) * indicates significant difference by T-Test (p<0.0045866, DF = 12-2). **H)** Same as G, but change in direction index values are indicated. *Left:* Difference is not significant (T-test, p<0.25972, DF = 12-2). *Right:* Difference is not significant (T-test, p<0.14439, DF = 12-2). Ns are animals (averages across all significantly-responding cells in each animal). The post-critical period animal was excluded in this analysis. RPI exhibited increases in inexperienced animals and in animals with low initial orientation index values.

While these data showed that the least mature animals acquired receptive fields that were more correlated with the phase-scrambled training stimulus, it remained possible that we were merely pushing the brain circuitry into an unnatural configuration that, while producing increased responses to the phase-scrambled training stimulus, was simply another allowable developmental configuration but not one that was truly instructed by the training stimulus. To understand how responses were altered relative to the full stimulus family, we projected the 10-d responses of these animals onto a reduced 2-d representation using principal component analysis (**Figure 7A**). In each case where we observed a significant training effect (full 95% range >0 in **Figure 6CE**), the mean responses of these animals moved closer to the training stimulus. Further, we performed a pairwise examination of the change in RPI for each stimulus compared to the training stimulus in these animals (**Figure 7B**). For most stimuli *(F,B,S1,S3,S4,CP1,CP2)*, the actual responses moved significantly closer to those of a hypothetical neuron that would respond optimally to the training stimulus. For other stimuli (S2, *S5, S6* when it was not the training stimulus) that were located near to the training stimuli (S4, S6) in 10-dimensional space (**Figure 7A**), the average tuning moved about equally close to hypothetical optimal responses for the reference stimulus and the training stimulus. In total, these results indicated that the neural responses were becoming more like those of hypothetical neurons optimally tuned to the training stimulus, as expected for an instructive influence.

**Figure 7.**
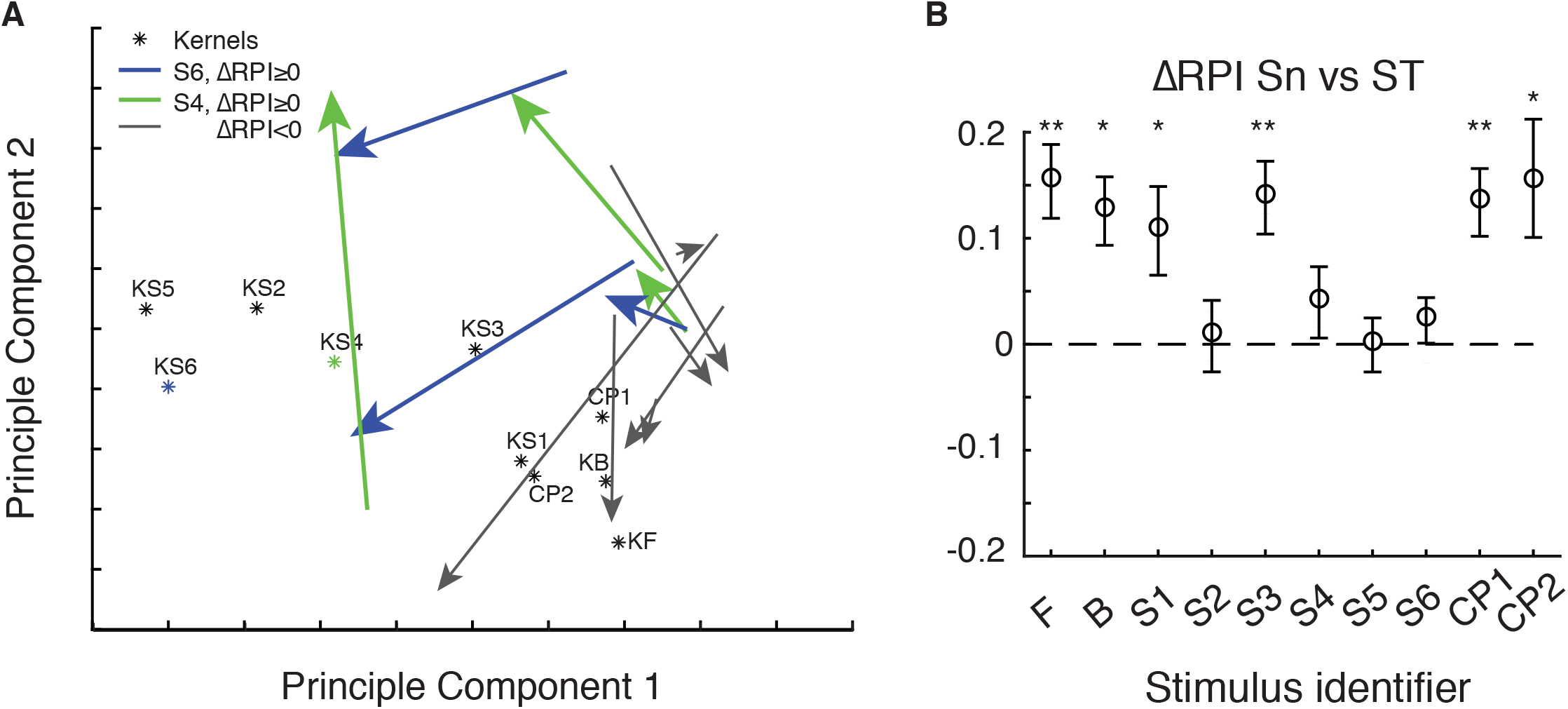
With phase-scrambled exposure, responses became more similar to those of ideal neurons selective to the exposed stimulus. **A)** Principle component projection from 10-dimensional space to 2-dimensional space of mean responses (for each animal) to the chosen set of 10 stimuli, before and after training, with vectors indicating the transition from the mean response before training to after training (arrow points at mean state after training). Responses of hypothetical neurons optimized for each stimulus (KF, S4, CP2 etc.) shown. Animals that exhibited significant ΔRPI (F vs. ST) are indicated (trained with S4 green, S6 blue). In this reduced view, average responses of significant animals moved closer to *KST*, while animals (8/8) that exhibited no significant effect moved to be near to *KF, KB, KCP1, KCP2* (typical V1 receptive fields). **B)** Change in RPI for significant animals with each stimulus used as a reference with the training stimulus *(Sn* vs. *ST*). For animals trained with S4 or S6, values of RPI (S4 vs. S4) or RPI (S6 vs. S6) were excluded from the average as it is 0 by definition. Error bars indicate SEM of the mean. * or ** indicates one-tailed T-test (* p<0.05, **p<0.005, DF = 6, DF = 3 when X is S4/S6) with mean > 0. Comparison for each RPI (X vs. ST) is single comparison evaluating only stimulus X. Changes in responses became more like a hypothetical neuron tuned to the training stimulus *KST* than stimuli F, B, S1, S3, S4, CP1, and CP2, and changes in responses remained equally close to stimuli S2, S5, and S6 (when S6 was *not* the training stimulus) on average. As responses changed in 10-dimensional space, they moved closer to *KST* for most stimuli while moving no closer or farther from KS2, KS5, and KS6. This is consistent with the idea that the training stimulus provided an instructive influence on receptive field properties in this early developmental period.

One may be concerned that the enhanced plasticity observed in the animals with least experience and weakest initial orientation index values could be artifactual coincidences if these animals also exhibited poorer quality responses. To address this possibility, we examined signal strength (that is, calcium response strength) and signal-to-noise ratios in **Supplementary Figure S4**, and found no correlation of signal strength or signal-to-noise ratio with experience or initial orientation index value. We therefore found no evidence that the quality of the responses differed across animals systematically by days of visual experience or initial orientation selectivity.

These results show that the spatiotemporal tuning of neurons was modified by visual experience provided to the premature cortex. However, all response properties were not malleable, indicating that the influence of premature or very early experience has limits. Orientation preference, for example, was not altered by this experience (**Supplementary Figure S5**), suggesting that either some features of the circuit are fixed even at our earliest point of examination, or that longer stimulation would be required to alter these properties. Nevertheless, the spatiotemporal response profile of these cells was modified through experience with a stimulus that was specific to the individual animal in a manner that was not possible just a few days later.

## Discussion

In this study, we tested the precise role of visual experience in the development of direction selectivity by exposing young ferrets to phase-scrambled gratings for several hours and asking if the cortical neurons could develop selective responses to such unusual stimuli. We found that in ferrets with no visual experience, V1 neurons developed selective responses to such irregular motion. In contrast, slightly more mature ferrets with 1-10 days of visual experience did not acquire increased selectivity to the phase-scrambled patterns, but instead exhibited a developmentally-typical increase in direction selectivity. We conclude that the influence of visual experience on the developing cortical circuit undergoes a transition – from instructive to permissive – right around the time of natural eye opening.

To our knowledge, this is the first time that cortical neurons have been induced to become selective to an irregular spatiotemporal stimulus through visual stimulation alone. In the disease amblyopia, a poor alignment of the 2 eyes or poor resolution in 1 of the 2 eyes causes a substantial drop in receptive field acuity/resolution and poor matching of receptive field properties across the 2 eyes^13^, which reflects degradation of receptive field structure but not the formation of a new, precise spatiotemporal receptive field. Other studies have imposed new receptive field structure, but have done so by pairing visual stimuli with external feedback control of visual^31^ or somatosensory^32^ cortex. The neurons in our study exhibited responses unlike those found in typically-developing animals in that they showed specific selectivity for an unnatural, phase-scrambled grating stimulus, which is a strong demonstration of instructive plasticity. This selectivity also differs from the interesting induction of sequence selectivity in visual cortex^33,34^ in that the selectivity introduced here is to a stimulus that is compact in space and time, with a cycle frequency of 4 Hz. The ability of cortex to acquire such unusual selectivity suggests that the cortex is particularly malleable in the face of activity in this very early window.

It is interesting that there are phenomenological differences among the forms of activity-driven plasticity for spatiotemporal scrambled stimuli (before or at eye opening), typical direction selectivity (first 2 weeks after eye opening) and ocular dominance plasticity (excluded until about 2 weeks after eye opening, then closes about 1 month after eye opening^35–38^). Functionally, it is useful for ocular dominance plasticity to be excluded around the time of eye opening, because one of the eyes may open earlier than the other. But spatiotemporal selectivity seems highly plastic immediately before and after eye opening. These phenomenological differences suggest that these forms of plasticity may be implemented by different mechanisms across development.

While we have shown that visual activity in the window from a few days before eye opening to just after eye opening drives strong plasticity in spatiotemporal selectivity, it remains unclear how exactly this plasticity is used by the developing animal under typical developmental conditions. Spontaneous activity, which is necessary for the development of orientation selectivity^9^, dominates in the week before eye opening^3,12,39^, and it may be the case that the patterns of this spontaneous activity “instruct” the formation of the cortical circuitry that reflects visual tuning parameters such as selectivity to smoothly moving stimuli. This spontaneous activity is sufficient for formation of orientation selectivity and the initial biases for direction angle preference because both still form in dark-reared animals^15,23^. In addition, very early visual experience through the closed lids drives visual activity^28,29^ and this activity, in addition to experience in the hours after eye opening, may instruct the development of smooth spatiotemporal receptive fields under typical conditions. Differences in the quality and patterning of activity that typically occurs in this early window may underlie species differences in functional architecture such as ocular dominance patches or receptive field parameters such as the fraction of cells that exhibit direction selectivity^40^.

We conclude that the influence of neural activity on the formation of visual circuits exhibits a sharp transition from instructive to permissive that occurs around the onset of natural visual experience. This conclusion builds on the prior knowledge that spontaneous activity before experience is necessary for proper development of visual circuits^3,9^ by suggesting that the quality and patterning of early activity sculpts the circuitry that supports the parameters of tuning such as spatiotemporal selectivity that are later revealed when selectivity is amplified through experience. After this transition, the net influence of activity-dependent circuit mechanisms must be qualitatively different, because a variety of patterns of neural activity drive the formation of typical smooth direction selectivity, with tuning parameters that cannot be greatly altered.

This developmental transition also mirrors a physiological transition observed in rats and in preterm humans, where flashes of light given before the typical onset of natural visual experience produce prolonged bursts of cortical activity, but these prolonged bursts fade around the onset of natural visual experience (P12 in rats, 36 gestational weeks in humans)^41^. The circuit changes underlying this transition are still unclear, but changes in cortico-thalamic loops and cortical inhibition may contribute^41,42^. Emerging research suggests that preterm humans exhibit higher rates of poor acuity later in life^43^ that cannot be explained by the acuity of the eyes^44^. The mechanisms underlying this poor acuity are not understood and may be varied. Brain damage could contribute^43^. But it is also possible that the premature cortex could be vulnerable to certain types of premature visual experience that could impact the formation of the initial brain circuitry. Future research on the influence of visual stimulation and neural activity on the premature brain may inform best practices for care of the earliest preterm infants.

## Methods Summary

Ferrets were anesthetized with ketamine and isoflurane (2% for surgery, 0.08-2% during imaging). GCaMP6s was introduced to the cortex with an AAV virus in a prior survival surgery. Changes in calcium fluorescence were monitored with a 2-photon microscope (Prairie Technologies) driven by a mode-locked laser (920nm, Mai Tai Deep See, Spectra-Physics). The training protocol consisted of 5s stimulation followed by 10s interstimulus interval; the protocol continued for 20min followed by 10min of no stimulation and this entire procedure was repeated for several hours. Stimulation and analysis were performed using custom software for Matlab (Mathworks). See **Supplementary Information** for details.

## Acknowledgements

We thank Jason Osik for help with experiments, Zoey Keeley and Daniel Shin for help with figure layout, and members of the Van Hooser lab for comments and feedback during the project and on the manuscript.

