## supplemental information for "An early phase of instructive plasticity in visual cortex before the typical onset of sensory experience"

### Supplementary Information

#### Contents Page

### Supplementary methods

#### Animal preparation

All experimental procedures were approved by the Brandeis University Animal Care and Use Committee and performed in compliance with National Institutes of Health guidelines.

***Animal source and housing.*** Ferrets (*Mustelo putorius furo*) were obtained from Marshall Bio-Resources. Litters of 4 or more kits arrived with a jill at around postnatal day (P) 10. Animals were housed in a room with timed lights (12 hours on, 12 hours off) in a custom stainless steel cage (60 cm x 60 cm x 35 cm) with a hammock and small toys.

***Virus injection.*** GCaMP6s was expressed in ferret V1 by transfection with the virus AAV2/9.Syn.GCaMP6s.WPRE.SV40[1] obtained from the UPenn Vector Core and later at AddGene (100843-AAV9). Detailed description of virus injection in young ferret kits has been published elsewhere[2]. Briefly, ferret kits (P19-20) were anesthetized with a ketamine-xylazine cocktail and a small craniotomy was made in the left hemisphere over visual cortex following sterilization of the scalp with Betadine and 70% isopropanol while the kit's head was held in a stereotactic device. Through a very small opening made in the dura, a glass injection micropipette (20-30uM tip diameter, beveled to ~22° using a Narishige Micropette Grinder EG-400) was inserted into the brain. With a volume injector (Nanoject II, Drummond Scientific), 1uL of virus solution (~$6.3 \times 10^{12}$ genomes/ml) was delivered over two different depths (150 and 500 um from the brain surface). The total volume was injected in small steps of 20–30nL with 10 to 15 sec rests between injections. A 10-14 day recovery period was allowed for GCaMP6s protein to express in visual cortical neurons before imaging experiments were carried out.

***Surgical preparation.*** The ferret was sedated with ketamine (20 $mg \cdot kg^{-1}$ im). Atropine (0.16–0.8 $mg \cdot kg^{-1}$ im) and dexamethasone (0.5 $mg \cdot kg^{-1}$ im) were administered to reduce bronchial and salivary secretion and to reduce inflammation, respectively. The animal was next anesthetized with a mixture of isoflurane, oxygen, and nitrous oxide through a mask and a tracheostomy was performed. The animal was then ventilated with 1.5–3% isoflurane in a 2:1 mixture of nitrous oxide and oxygen. A cannula was inserted into the intraperitoneal (ip) cavity for delivery of neuromuscular blockers and Ringer solution (3 $ml \cdot kg^{-1} \cdot h^{-1}$), and the animal was inserted in a custom stereotaxic frame that did not obstruct vision. All wound margins were infused with bupivacaine. Silicone oil was placed on the eyes to prevent corneal damage. Before imaging commenced, the ferret was paralyzed with the neuromuscular blocker gallamine triethiodide (10–30 $mg \cdot kg^{-1} \cdot h^{-1}$) through the ip cannula to suppress spontaneous eye movements, and the nitrous oxide-oxygen mixture was adjusted to 1:1. The animal's ECG was continuously monitored to ensure adequate anesthesia, and the percentage of isoflurane was increased if the ECG indicated any distress. Body temperature was maintained at 37°C.

While the animal's head was held in the stereotaxic frame, a wire mesh was attached to the skull with VetBond (3M) glue to provide an anchor point for dental acrylic, and a stainless steel head plate with a 1 cm opening was cemented to the skull above the virus injection site[3].

The head plate was secured to the skull with dental acrylic fortified with VetBond glue and cured with Zip Kicker (ZAP). A small opening in the cranium was drilled through the head plate opening with a dental drill (Medidenta). The dura above the injection site was removed over a 1mm x 1mm area for imaging. Warm agarose (3% in 0.1M phosphate buffered saline) was poured over the opening and a 5 mm circular coverslip (Warner Instruments) was gently pressed into the solidifying agarose such that the coverslip rim touched the bone around the craniotomy and small amounts of agarose oozed out on top of the coverslip rim to secure it in place.

#### Visual stimulation

Visual stimuli were created in MATLAB with the Psychophysics Toolbox[4-6] on a Macintosh Pro running OS X (10.6, Snow Leopard) and displayed on a Dell monitor 1704FPVt (40 cm viewing distance) and gamma-corrected with a Spyder 3 Express. All stimuli were sinusoidal gratings presented on the full screen at a spatial frequency of 0.8 cycles per degree and a temporal frequency of 2Hz (discretized in the case of phase-scrambled sequences). All gratings were displayed as 10 repeating cycles for a duration of 5 seconds, with an interstimulus interval of 5 seconds. During training, stimuli were displayed 5 seconds on, 10 seconds off, in bouts of 20 minutes, with 10 minutes of rest (gray screen) after each 20 minute bout.

#### 2-photon calcium imaging

Cells in V1 were imaged with a 2-photon microscope (Ultima IV, Prairie Technologies, Madison, WI) using 920 nm laser light (Mai Tai Deep See, Spectra Physics) and a 16x saline-immersed objective (16x, 0.8NA Nikon) with total output power < 50 mW. To shield the light of the stimulus monitor from the microscope, the objective and head plate were covered with a custom-sewn sleeve made from light-tight fabric (ThorLabs) and elastic (Michaels Stores Inc.), and augmented as needed with light-tight tape (ThorLabs)[3]. Light block quality was assessed by setting the photomultiplier gain to maximum, shuttering the laser, and scanning while turning the monitor on and off to verify that no modulation could be observed. During visual stimulation, 512x512 pixel image frames were acquired continuously every 1.3-1.8 s, and 8 repetitions of each stimulus were recorded.

Cells were imaged in layer 2/3 at depths ranging from 100 µm to 300 µm. In all cases, we examined the cytoarchitecture at the surface and focused downward to be sure we were lower than cortical layer 1. Before training, images were typically acquired only at a single depth to avoid providing too much experience to the animals during the measurements (each measurement was 20 minutes). After the training, images were typically acquired at more than 1 depth, and these recordings were displaced in Z by at least 40 µm to avoid recording the same cells twice. All recordings were carefully examined to ensure stability in the Z axis, and any recordings showing significant Z drift were discarded.

#### 2-photon imaging analysis

Images were analyzed with custom software written in Matlab and C. Cells were identified by the experimenter, who selected regions of interest for GCaMP6s with new custom software extensions of our 2-photon analysis tools (http://github.com/VH-Lab/vhlab-twophoton-matlab). The experimenter watched movies of the responses in order to detect neurons that only exhibited strong fluorescence when active, and could select regions-of-interest in any video frame. Small horizontal drift over time was corrected manually with a custom software tool that allowed the user to track and align landmarks in the movies over time (correlation-based methods, which work well for images with stable backgrounds such as those obtained with OGB-Bapta-1 AM[7,8], did not work well for GCaMP6s). For each frame, the intensity value of each cell was computed by averaging all pixels in a circle of radius ~10 µm that was centered on the soma. These circular regions of interest were smaller than the soma itself to reduce possible intrusion of signals from the surrounding neuropil. Sometimes, the same cells could be identified across subsequent imaging sessions in the same animal (hours apart in time, see [8,9] for methods), but often this was not the case because cells were dim when they were not active. (The experiment in **Figure 4** was a case where we could make several cell correspondences before and after training.) We made no attempt to specifically report summary data in cells that were recorded at multiple time points in this paper, and simply examine all cells recorded "before" several hours of stimulus exposure and compare these responses to all cells recorded "after".

#### Software

The Matlab code used to run stimuli and perform analyses are in the Van Hooser lab GitHub distributions (see https://github.com/VH-Lab/vhlab_vhtools/wiki/Installation for installation of all packages).

#### Stimulus design

Our goal was to develop a stimulus set that could be delivered at a fixed orientation, in order to drive cortical neurons, but that would exhibit a variety of spatiotemporal patterns within that orientation. We designed sinusoidal gratings that moved in irregular temporal patterns. For this purpose, we discretized one full temporal cycle of a grating into 8 phase steps. In this scheme, one cycle of regular forward or backward motion was created by moving a sine-wave grating following an ordered sequence of 8 phase steps ([1 2 3 4 5 6 7 8] or [8 7 6 5 4 3 2 1]), such that on the 9$^{th}$ step of the temporal sequence the grating arrived at the starting spatial phase. To create irregular motion, the sine-wave grating was moved following a scrambled sequence of 8 steps (e.g., [7 4 8 3 1 6 2 5]) to complete one full cycle. Any such 8-step temporal sequence, regular or scrambled, was repeated 10 times to create a 5 second stimulus. Each phase step was shown for 1/(2Hz*8) = 0.0625 s to reflect a repetition rate of 2 Hz.

In determining the requisite number of phase steps to define a full grating cycle, we considered 3 factors.  First, the duration of each phase step on the screen had to be long enough for the visual system to respond to it. If we used a very high number of phase steps, then the temporal energy of a scrambled phase sequence would be very high and the visual system would not respond. Second, the number of phase steps had to be large enough to generate sufficiently distinct stimuli; if we had chosen 4, for example, then only 6 sequences were possible (including forward and backward). Third, the number also had to be small enough so that the total number of possible combinations was within practical limits of analysis. We chose 8 phase steps and 2Hz animation to be a compromise among all of these factors.

A grand total of 40,320 (= 8!) sequences can be created out of 8 steps with no repeats allowed. However, when a sequence is repeated multiple times, as in showing multiple cycles of the grating, the sequence should be considered in a circular space. Effectively, sequence [1 2 3 4 5 6 7 8] is the same as sequence [2 3 4 5 6 7 8 1]. In the circular space, each sequence will have 8 variants that differ only in the starting phase. Thus, the total number of *unique*, non-phase-shifted sequences made of [1:8] reduces to 5040 (= 40320/8). Out of these, 2 sequences represent regular motion ([1 2 3 4 5 6 7 8] = forward; [8 7 6 5 4 3 2 1] = reverse), while the rest represent irregular motion.

Before choosing a small number of sequences out of the set of 5040 for experiments, we quantified the degree to which each sequence differed from smooth motion, by calculating 3 separate metrics. First, we calculated the correlation between the sinusoidal phase shifts for all 5040 unique sequences and forward and backward motion (**Fig. 1C**). Correlation was calculated for all possible phase offsets between the 2 sequences in order to find the value with the highest correlation. For example, sequence [3 2 4 5 6 7 8 1] would be rotated to [1 3 2 4 5 6 7 8] when calculating correlation with [1 2 3 4 5 6 7 8]. Note that the calculation is a sum of products of 2 sinewaves at each phase step ($\sin(2\pi((step_1-1)/8)x) * \sin(2\pi((step_2-1)/8)x)$), where x is the spatial variable. We calculated phase with x = [0 1 2 … 10] / 10. For each sequence, the correlations were calculated against forward and backward smooth sequences separately (**Fig. 1C**), and then the two values were averaged to obtain an average correlation to smooth motion Supplementary Figure 1C). Second, we calculated the summed total phase interval for each sequence (Supplementary Figure 1A, C, D). For sequences representing smoother motion (e.g., [1 2 3 4 5 6 7 8]), the phase intervals or steps between adjacent phases are small, thereby yielding a smaller summed total phase interval value. In contrast, a sequence representing higher degree of irregularity (e.g., [8 3 6 2 7 4 1 5]) would contain larger phase steps and therefore yield a larger summed total phase interval value. Third, we computed the 2-D Fourier spectra for each sequence (Supplementary Figure 1B) and then calculated the total motion energy content for the sequence by summing the absolute value of all Fourier coefficients excepting the 0 temporal frequency value (static grating).

All 3 measures captured overlapping but slightly different aspects of the irregularity of the sequences. Our goal was to choose a small set of sequences for experiments that covered different levels of irregularity. To do so, we first generated various scatter plots of the 3 irregularity measures against each other (**Figure 1C, Supplementary Figure 1CD**). From these plots, it is generally evident that sequences with higher summed phase intervals also tended to have lower correlation to smooth motion sequences and higher total motion energy, thereby representing more irregular motion.  From these scatter plots, we hand-picked a set of 6 sequences ("scrambled"; S1-6) such that some of them fell on the higher end of dissimilarity to smooth motion and some fell in between.  In addition, we included 2 classic counter-phase grating stimuli sequences that represent the sum of forward and backward smooth motion (CP1 and CP2), which are not part of the set of 5040 sequences analyzed.  We deliberately chose not to include any sequence too similar to smooth motion sequences, because any positive effect of training with such stimuli could simply be the result of smooth motion training.

In **Supplementary Figure 2** we show examples of the 2-D Fourier spectra of some of the chosen 10 sequences. The smooth gratings have energy at a single spatial and temporal frequency (positive and negative), while the scrambled gratings exhibit spatial and temporal energy at several spatial/temporal frequency values and show power at higher temporal frequencies than the smooth versions. This results in these sequences having greater total motion energy.

#### Analyses of responses and statistics

The response to each stimulus was calculated as ($F_{stim}$-$F_{base}$)/$F_{base}$, where $F_{stim}$ is the average response inside the region-of-interest during each frame when the stimulus was on, and $F_{base}$ is the average response during the final 3 seconds of the interstimulus period before stimulus onset. In the youngest animal, aged P29, responses continued for up to 2 seconds after the stimulus turned off, and the response was averaged in an interval [0 7] seconds post-stimulus as opposed to [0 5] seconds for most experiments.

Cells were categorized as "visually responsive" if an ANOVA test over all stimuli and the blank stimulus yielded p<0.05. Cells/regions-of-interest that were not visually responsive were ignored.

The orientation selectivity index for visually-responsive cells was assessed using 1 minus the circular variance, calculated in orientation space[10]:

$$1 - CirVar = \left| \frac{\sum_k R(\theta_k)\exp(2i\theta_k)}{\sum_k R(\theta_k)} \right|,$$

where $R(\theta_k)$ is the response to angle $\theta_k$ with the response to the blank stimulus subtracted.

Cells were categorized as exhibiting significant orientation tuning if responses to drifting gratings of varying orientations passed Hotelling's $t^2$-test with p<0.05[8,11].

If cells were orientation-selective, then responses were fit with a double gaussian function[11-13]:

$$R(\theta) = C + R_p e^{-\frac{\mathrm{ang}_{\mathrm{dir}}(\theta-\theta_{pref})^2}{2\sigma^2}} + R_n e^{-\frac{\mathrm{ang}_{\mathrm{dir}}(\theta+180-\theta_{pref})^2}{2\sigma^2}},$$

where $C$ is a constant offset, $\theta_{pref}$ is the preferred orientation, $R_p$ is the above-offset response to the preferred direction, $R_n$ is the above-offset response to the null direction, and $\text{ang}_{\text{dir}}(x) = \min(x, x-360, x+360)$ wraps angular difference values onto the interval 0° to 180°, and $\sigma$ is a tuning width parameter. The tuning width (half-width at half-height) is equal to $\sqrt{\log 4}\sigma$ (half-width at half height)[12]. For these cells, the direction index value was calculated as DI = (Rp-max(Rn,0))/Rp. The *max* function constrains DI to be at most 1, following [8].

The Selectivity Index (SI) for stimulus *n* given responses to stimuli *[F, B, S1,…,CP2]* was computed as:

$$SI(n) = \frac{R(Sn)}{R(F)+R(B)+R(S1)\cdots R(S6)+R(CP1)+R(CP2)}$$

The Selectivity Index could, in principle, vary from 0 (no response to *Sn*) to 1 (exclusive response to *Sn*), but in practice it tended to peak at lower values (peaks of about 0.2-0.3) because many of the chosen 10 stimuli are correlated with each other. For example, a cell that responded to forward motion would be very likely to respond to the counterphase stimuli CP1 and CP2, because stimuli CP1 and CP2 are mathematically equal to the sum of gratings drifting past one another: that is, they are mathematically equal to the sum of stimuli F and B.

We developed a second measure that considered the responses to all 10 of the stimuli that we termed the Response Projection Index (*RPI*). We can imagine neural kernels (*KX*) that would give a maximal response to each stimulus (*X*), as shown in **Fig. 1EF**, and we can compute the responses of these kernels to each of the 10 stimuli. The kernels were taken to be sinewave X-T profiles that matched the structure of the stimulus, except reversed in time. Responses of a kernel to another stimulus was computed by taking the correlation (with optimal phase alignment) between the stimulus and the kernel, and normalizing the kernel's response to its preferred stimulus as 1. The RPI describes how close the response of a measured neuron, in the 10-dimensional response space, is to the responses that would be expected from one ideal kernel (*KXi*) relative to another ideal kernel (*KXj*). The *RPI* is therefore a relative measure, and requires 2 referent kernels in addition to the measured responses. For example, a neuron that gives responses that are identical to *KF* would have an *RPI(KF vs KB)* value of -1, indicating that the response is close to *KF*, whereas a neuron that gives responses that are identical to *KB* would have an *RPI(KF vs KB)* value of +1. Stimulus *S6* is uncorrelated with *F* and *B*, and *KS6* has an *RPI(KF vs KB)* value of 0.

Given a 10-dimensional vector of responses **R** of a cell and $KX_i$ and $KX_j$, we calculated

$$D1 = \left\| R - (R \bullet u_1)u_1 \right\|, u_1 = R(KS_i) / \left\| R(KS_i) \right\|$$
$$D2 = \left\| R - (R \bullet u_2)u_2 \right\|, u_2 = R(KS_j) / \left\| R(KS_j) \right\|$$

where ||x|| is the Euclidean norm of the vector ***x***, and then

RPI ($KS_i$ vs. $KS_j$) = (D1-D2)/(D1+D2).

***Bootstrap confidence intervals***: To examine whether a change in RPI was greater than 0 and to put confidence intervals on this distance, we performed a bootstrap difference analysis for each animal. Assume the population of neurons examined before training was N and the population examined after training was M. We then created 10,000 bootstrap simulations where we drew N neurons from the before population (with replacement), and M neurons from the after population (with replacement), and calculated the difference Δ. We then had a distribution of 10,000 values of Δ. We took the mean of this distribution as the mean difference, and reported the 5% and 95% values of this distribution Δ as the low and high confidence intervals, respectively. If the low confidence value was greater than 0, then the difference in RPI was said to be significant.

To examine the significance of the correlation between days of experience or initial orientation selectivity index values and ΔRPI or ΔDI, we calculated the p-value of the correlation coefficient using the Matlab function **`corrcoef`**. T-tests in Figure 6GH were performed with the Matlab function **ttest2**.

#### Statistics table

| Result | p-value | Test employed | Degrees of freedom | Additional information |
|---|---|---|---|---|
| **Non-significant** correlation between days of visual experience and ΔRPI (KF vs. KST) (**Fig. 6C**) | p<0.1650 | **corrcoef** | 12-2 | Post-critical period animal excluded. Empirical correlation coefficient was **negative**. |
| **Significant** correlation between initial orientation selectivity and ΔRPI (KF vs. KST) (**Fig. 6E**) | p<0.009 | **corrcoef** | 12-2 | Post-critical period animal excluded. Empirical correlation coefficient was **negative**. |
| **Non-significant** correlation between days of visual experience and ΔDI (**Fig. 6D**) | p<0.0678 | **corrcoef** | 12-2 | Post-critical period animal excluded. Empirical correlation coefficient was **positive**. |

| | | | | |
|---|---|---|---|---|
| **Non-significant** correlation between initial orientation selectivity and ΔDI (**Fig. 6F)** | p<0.1524 | **corrcoef** | 12-2 | Post-critical period animal excluded. Empirical correlation coefficient was **positive**. |
| **Significant** difference between inexperienced and experienced animals (**Fig 6G, left**) | p<0.020241 | **ttest2** | 12-2 | |
| **Significant** difference between inexperienced and experienced animals (**Fig 6G, right**) | p<0.0045866 | **ttest2** | 12-2 | |
| **Non-significant** difference between inexperienced and experienced animals (**Fig 6G, left**) | p<0.25972 | **ttest2** | 12-2 | |
| **Non-significant** difference between inexperienced and experienced animals (**Fig 6G, left**) | p<0.14439 | **ttest2** | 12-2 | |

#### Resource table

| Reagent type | Designation | Source or reference | Identifiers | Additional information |
|---|---|---|---|---|
| Software | Matlab | The MathWorks, Natick, MA | RRID: SCR_001622 | |
| Software | GitHub | GitHub | RRID: SCR_002630 | |
| Software | Psychophysics Toolbox | Psychtoolbox.org | RRID: SCR_002881 | |
| Virus | AAV2/9.Syn.GCaMP6s. WPRE.SV40 | UPenn Vector Core (previously) and now AddGene 100843-AAV9 | Plasmid is RRID:Addgene_100843 | Gift from the GENIE Project & Douglas Kim |
| Hardware and software | Spyder Express 3 | Datacolor | | |
| Microscope | Ultima IV 2-photon microscope | Prairie Technologies (now Bruker) | | |
| Laser | Mai Tai HP Deep See | Spectra-Physics / Newport | | |
| Objective | Nikon 16x | Nikon | CFI75 LWD 16X W | 0.8 NA, 3.0mm WD |

| | | | |
|---|---|---|---|
| Precision injector | Nanoject II | Drummond Scientific | |
| Objective sleeve fabric | Black nylon, polyurethane-coated fabric | ThorLabs | BK5 |
| Light-block tape | Black masking tape | ThorLabs | T743-1.0, T743-2.0 |
| Animals | Ferrets | Marshall Bio-resources | “Conventional” colony |
| Beveler | Micropipette grinder | Narishige | EG-400 |

### Supplementary Figures and Observations

#### S1: Quantitative analysis of phase-scrambled sequences

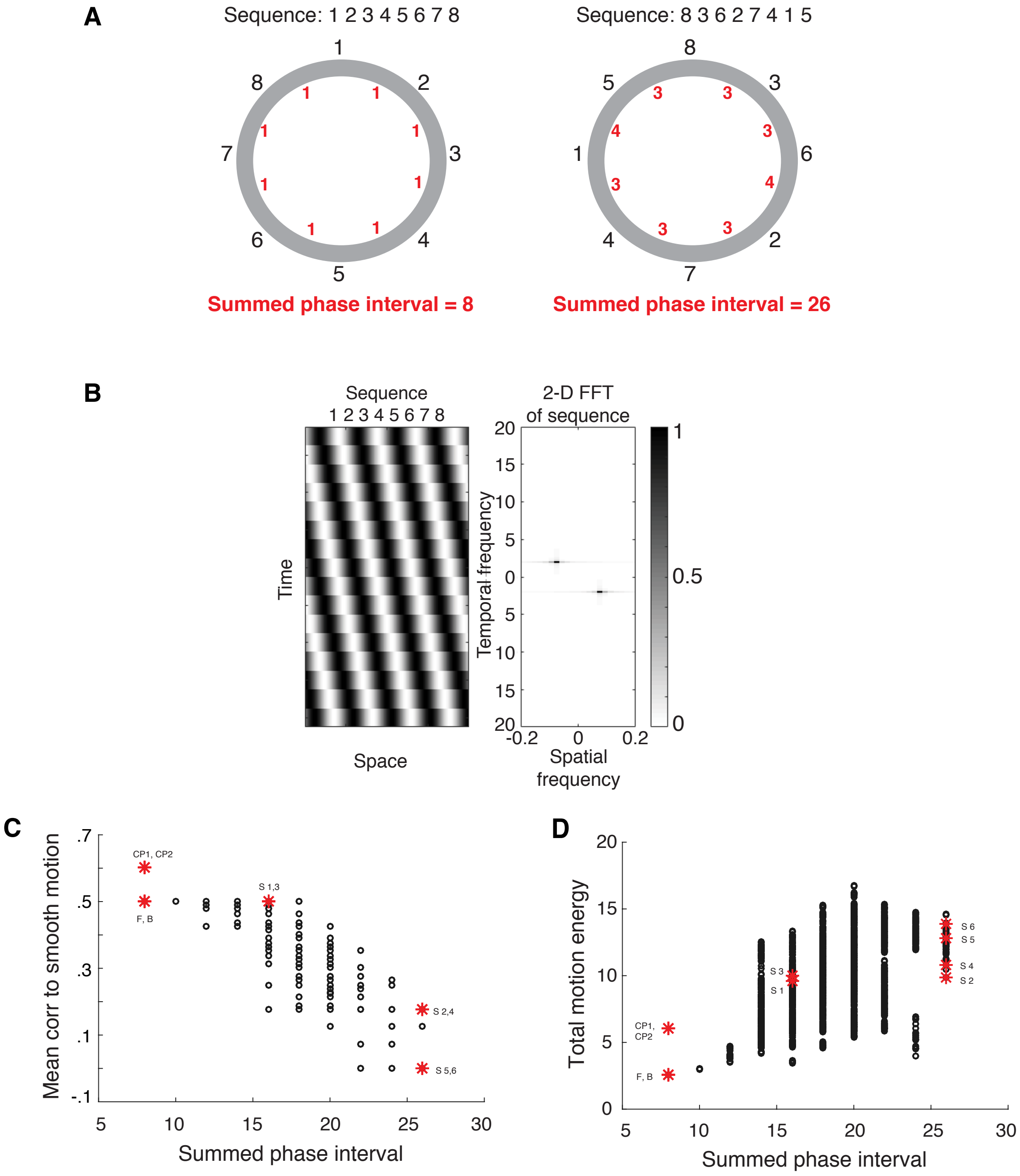

**Supplementary Figure S1. Quantitative analysis of phase-scrambled sequences. A)** Summed total phase interval. 2 example sequences built out of 8 phase steps are shown in circular space. The phases are shown in black digits, while the absolute value of the differences between two consecutive phases are depicted in red digits. The sum of all the phase intervals are shown below each sequence. **B)** 2-D Fourier spectrum of sequence [1 2 3 4 5 6 7 8]. On the left, 2 cycles of the grating stimulus are shown in space-time, moving following the sequence [1 2 3 4 5 6 7 8]. On the right is shown the corresponding 2-D Fourier spectrum, where each pixel represents the combination of specific spatial and temporal frequencies and the darkness of the pixel represents the normalized Fourier coefficient at those frequency combinations. To calculate total motion energy, all Fourier coefficients within two diagonal quadrants of the 2-D spectrum were summed, except for the pixels corresponding to temporal frequency 0, which represents static grating. **C)** Scatter plot of average correlation to smooth motion against summed phase interval, for all 5040 sequences. The apparent low number of points on the plot is due to very high overlap of these metrics between many sequences. **D)** Scatter plot of total motion energy against summed phase interval, for all 5040 sequences. In both C and D, the locations of the chosen 10 sequences (F, B, S1-6, CP1, CP2) on the space are marked with red asterisks.

#### S2: Spatiotemporal Fourier spectrum of stimuli

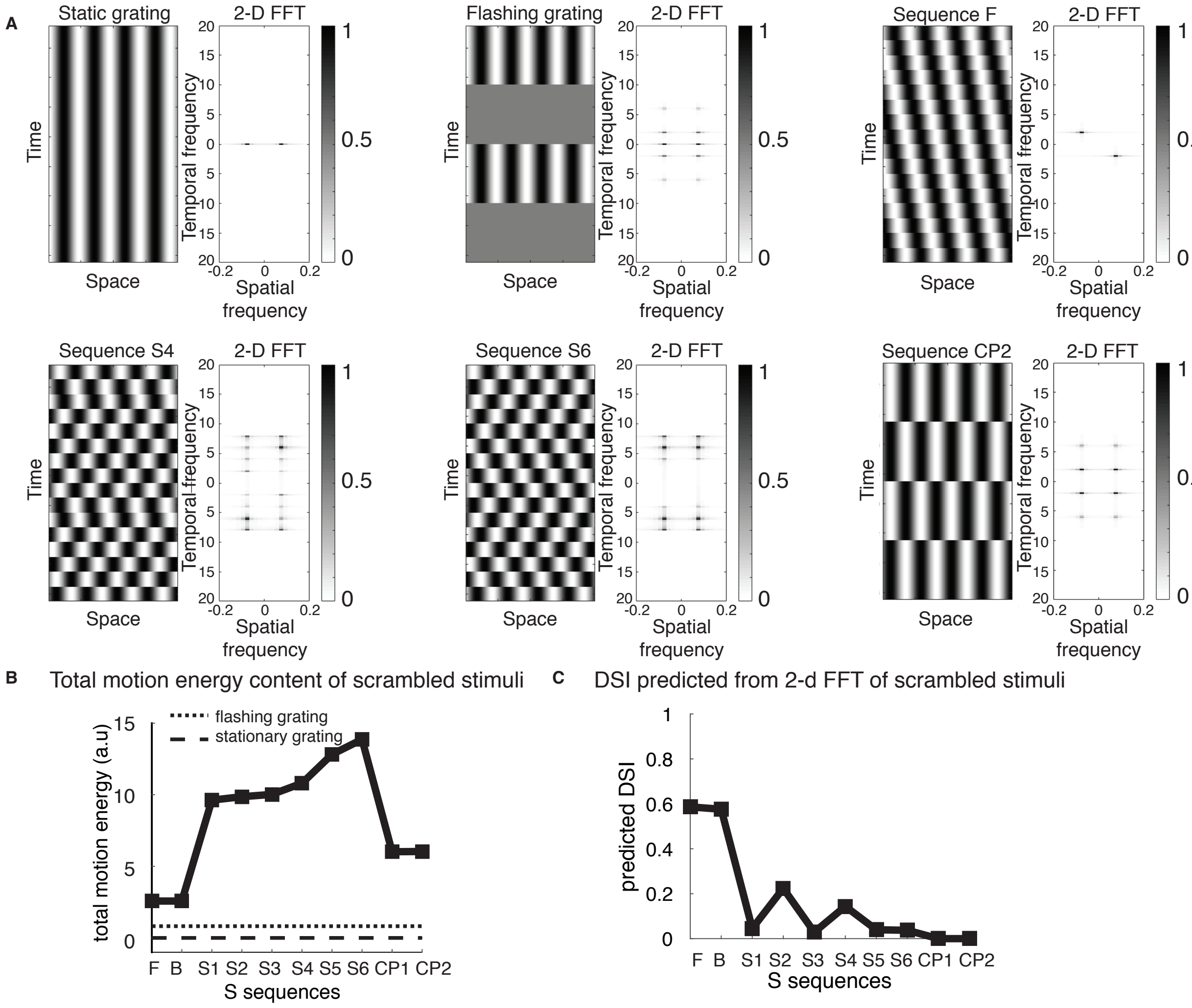

**Supplementary Figure S2. Spatiotemporal Fourier spectrum of stimuli. A)** 2 cycles of each grating stimulus sequence indicated and the corresponding 2-D Fourier spectrum. A static grating exhibits energy at a single spatial frequency (positive and negative values). A flashing grating exhibits energy at more spatial/temporal frequency values. A smoothly moving grating (F) exhibits energy at a single pair of spatial/temporal frequency values. The phase-scrambled sequences S4 and S6 exhibit energy at a variety of the higher harmonics of the underlying temporal frequency (2Hz), and therefore has energy at more temporal frequencies. **B)** Total motion energy (sum of absolute value of Fourier coefficients excepting the 0 temporal frequency value) of the chosen 10 sequences. **C)** Direction selectivity estimated by convolving each sequence with forward and backward motion and computing (Response_Pref – Response_Opposite) / Response_Pref.

#### S3: Changes in stimulus selectivity for all animals in the study

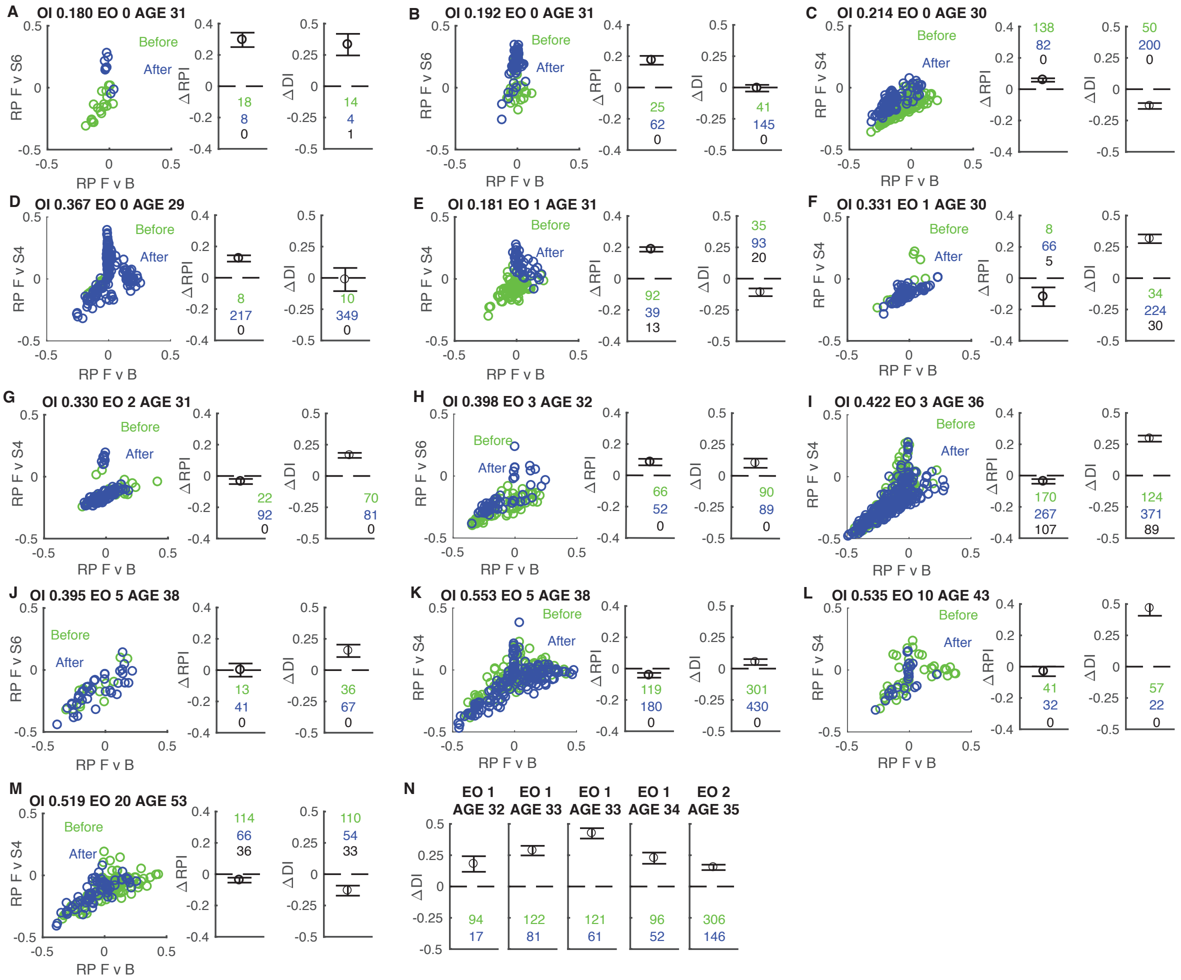

**Supplementary Figure S3**. **Changes in stimulus selectivity for all animals in the study.** Animals are ordered by days of visual experience (EO) and initial orientation index. **A-M**) *Left*: Response Projection Index (RPI) for *F* vs *B* (X axis) and *F* vs the trained stimulus (*S4* or *S6*, depending) (Y axis) for cells measured before (green) and after (blue) 6 hours of experience with the training stimulus. Days after eye opening (EO) are noted, as is the animal's postnatal age in days. *Middle*: Estimated difference in RPI (*F* vs. trained stimulus) before and after experience (95% confidence intervals). Cell N before (green) after (blue) and number of cells tracked across the imaging sessions (black) indicated. *Right*: Estimated difference in DI (*F* vs. trained stimulus) before and after experience (95% confidence intervals). Cell N before (green) after (blue) and number of cells tracked across the imaging sessions (black) indicated. We noted that, on average, animals with fewer days of natural visual experience were more likely to exhibit increased selectivity to an arbitrary stimulus that was presented for 6 hours, whereas animals with several days of visual experience were less likely to exhibit increased selectivity to this stimulus. Instead, animals that had 1-10 days of visual experience often exhibited increases in basic direction selectivity rather than selectivity for the phase-scrambled stimulus. **N)** Change in direction selectivity in 5 experiments from Li/Van Hooser et al. (2008). All animals exhibited robust increases in direction selectivity when bidirectional moving stimuli (3-6 hours) were used as the training stimulus.

**S4: Lack of significant correlation of days of visual experience or initial orientation selectivity with signal-to-noise or signal strength**

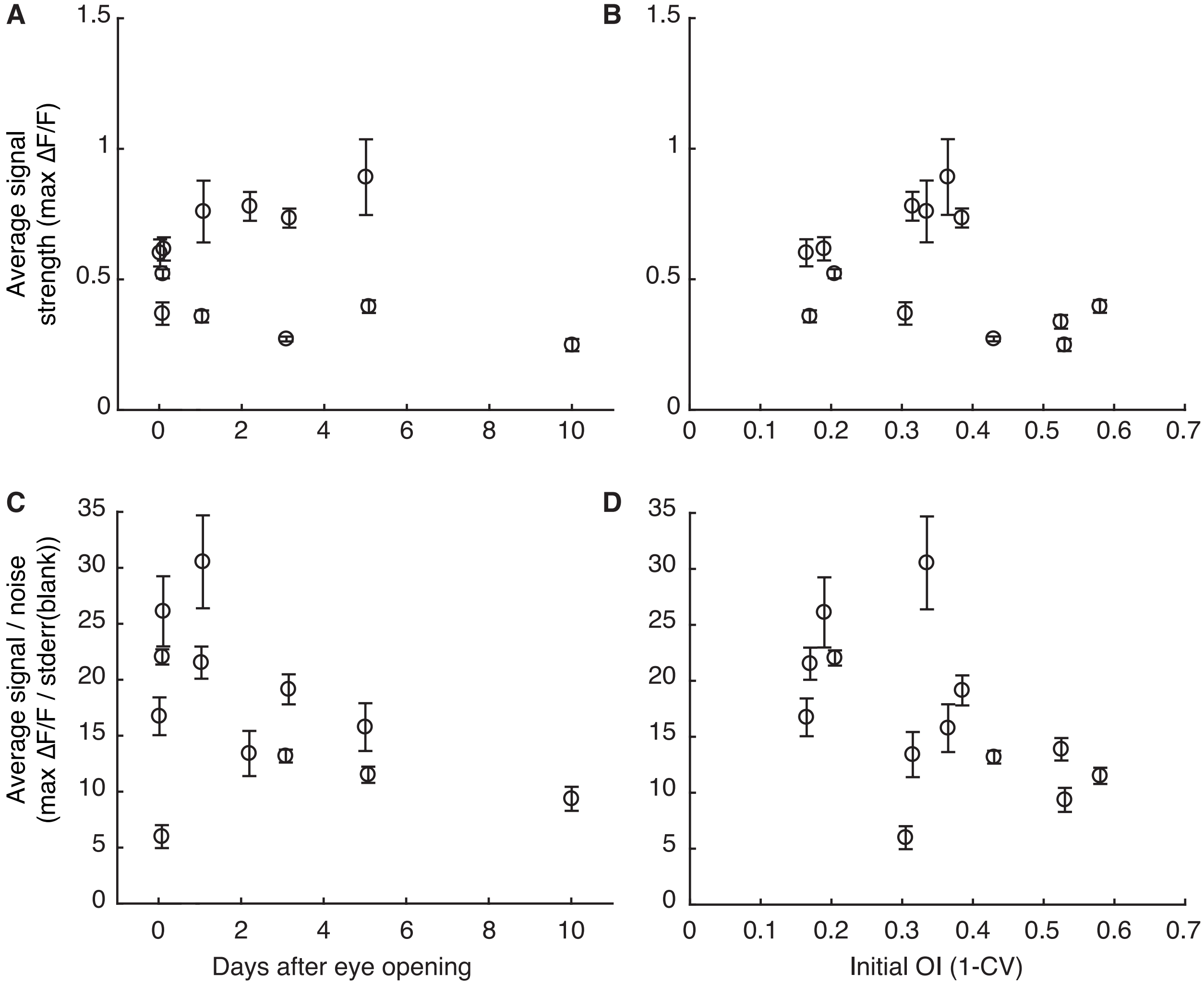

**Supplementary Figure S4**. **Lack of significant correlation of days of visual experience or initial orientation selectivity with signal-to-noise or signal strength.** Initial average signal strength (mean ΔF/F response to the preferred stimulus after blank stimulus response subtraction) plotted against **A)** days after eye opening (Correlation test: p=0.4516, DF = 12-2) and **B)** initial orientation index (Correlation test: P=0.4081, DF = 12-2). Signal to noise ratio defined as signal strength (defined as in **A** and **B**) divided by the standard error of the mean of the response to the blank stimulus plotted against **C)** days after eye opening (Correlation test: P=0.1334, DF = 12-2) and **D)** against initial orientation index (Correlation test: P=0.0939). The average value of these quantities was computed across all cells for each animal, and the mean and standard error of the mean are shown as each data point on the graph. These results indicate that the results of the study cannot reflect very poor or noisy responses in the youngest or most immature animals.

#### S5: No changes in orientation preferences over training

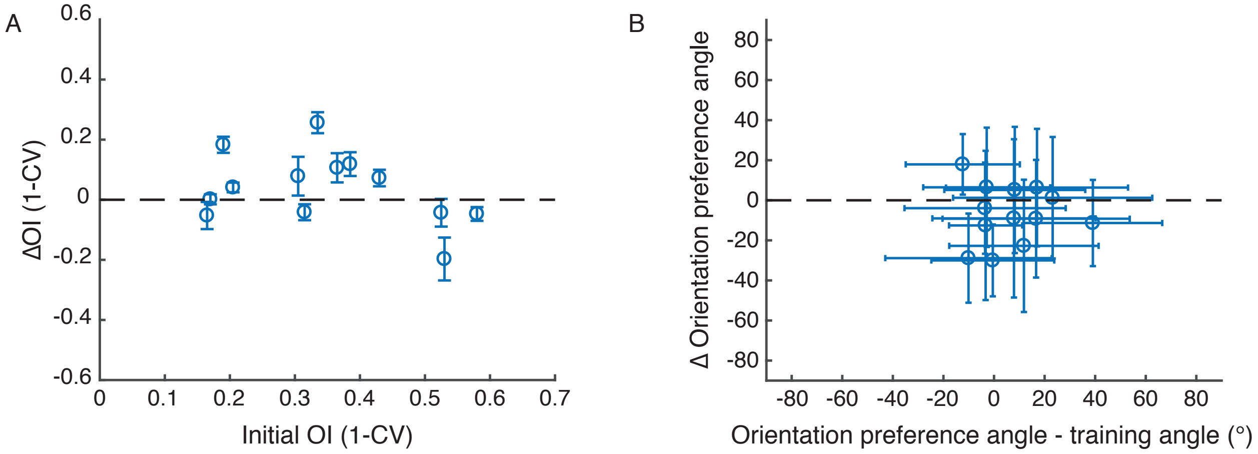

**Supplementary Figure S5**. **No changes in orientation preferences over training. A)** Changes in orientation selectivity index values before and after training as a function of initial orientation selectivity. On average, orientation selectivity index values increased slightly after training, but not consistently from case to case. Mean and bootstrap standard error is shown for each animal. **B)** Changes in orientation preference values for each animal with respect to initial orientation preference angle, with circular standard deviation shown (in degrees). The orientation preference angle was rotated so that the training orientation was 0. If a strong change in orientation preference were evident, the line should have a negative slope (so that negative rotated orientations would be increased, and positive rotated orientations would be decreased). Instead, there is no correlation. Correlation test: P=0.987, degrees of freedom = 13-2.

**Supplementary videos:**

F – sequence [1 2 3 4 5 6 7 8] “forward”

B – sequence [8 7 6 5 4 3 2 1] “backward”

S1 – sequence [8 1 2 7 4 5 6 3] “scrambled 1”

S2 – sequence [8 4 7 3 6 2 5 1]

S3 – sequence [1 4 3 2 5 8 7 6]

S4 – sequence [1 5 2 6 3 7 8 4]

S5 – sequence [1 6 3 8 4 7 2 5]

S6 – sequence [8 3 6 2 7 4 1 5]

CP1 – sequence [3 3 3 3 7 7 7 7] “counterphase 1”

CP2 – sequence [1 1 1 1 5 5 5 5] “counterphase 2”
